# On the role of different age groups during pertussis epidemics in California, 2010 and 2014

**DOI:** 10.1101/405076

**Authors:** Ayesha Mahmud, Marc Lipsitch, Edward Goldstein

**Affiliations:** Center for Communicable Disease Dynamics, Department of Epidemiology, Harvard T.H. Chan School of Public Health, 677 Huntington Ave., Boston, MA 02115, USA; Department of Immunology and Infectious Disease, Harvard T.H. Chan School of Public Health, 677 Huntington Ave., Boston, MA 02115, USA

## Abstract

**Background:** There is limited information on the roles of different age groups in propagating pertussis outbreaks, and the temporal changes in those roles since the introduction of acellular pertussis vaccines.

**Methods:** The relative roles of different age groups in propagating the 2010 and the 2014 pertussis epidemics in California were evaluated using the RR statistic that measures the change in the group’s proportion among all detected cases before-vs.-after the epidemic peak.

**Results:** For the 2010-11 epidemic, evidence for a predominant transmission age group was weak, with the largest RR estimates being 1.26(95%CI (1.08,1.46)) (aged 11-13y); 1.19(1.01,1.4) (aged 9-10y); 1.17(0.86,1.59) (aged 14-15y); 1.12(0.86,1.46) (aged 16-19y); and 1.1(0.89,1.36) (aged 7-8y). The 2014 epidemic showed a strong signal of the role of older adolescents, with the highest RR estimate being in those aged 14-15y (RR=1.83(1.61,2.07)), followed by adolescents aged 16-19y (RR=1.41(1.24,1.61)) and 11-13y (RR=1.26(1.12,1.41)), with lower RR estimates in other age groups.

**Conclusions:** As the time following introduction of acellular pertussis vaccines in California progressed, older adolescents played an increasing role in transmission during the major pertussis outbreaks. Booster pertussis vaccination for older adolescents with vaccines effective against pertussis transmission should be considered with the aim of mitigating future pertussis epidemics in the community.

## Introduction

After decades of low pertussis activity in the US following the introduction of whole-cell pertussis vaccines in the late 1940s, major outbreaks took place during 2004-2005, 2010, 2012 and 2014 [1]. Several factors behind the increase in reported pertussis incidence were proposed [2,3], such as the introduction of acellular pertussis vaccines and the reduction in protection compared to the receipt of whole-cell vaccines [4,5], waning effectiveness of acellular vaccines [6-10], improved testing and reporting [11], and the possible impact of genetic changes to *B. pertussis* [12,11]. A related question in pertussis epidemiology is the relative importance of individuals in different population groups in propagating pertussis outbreaks, and the potential impact of vaccination on the spread of pertussis in the community, including the disease burden in infants [13]. An important aspect of this issue if the fact that, as the time from the introduction of acellular vaccines in different places increases, children of increasingly older ages are covered entirely by the acellular pertussis vaccination series. In light of the evidence of decreased protection associated with receipt of acellular pertussis vaccines alone compared to receipt of some whole cell pertussis vaccination [4,5], the role of older children during pertussis outbreaks is generally expected to increase with time. Furthermore, the efficacy of the Tdap vaccine, usually administered around the age of 11 years [14] wanes with time since vaccine administration [8,10]. Additionally, pertussis vaccine effectiveness against infection and transmission to others can be lower than the effectiveness against symptomatic disease episodes [15], and the difference in effectiveness against infection vs. detected symptomatic disease is potentially more pronounced for Tdap compared to DTaP [16], with a survey of evidence about herd immunity effects of DTaP given in [17]. All of this suggests that older adolescents may potentially play a prominent role in propagating the more recent, as well as future pertussis epidemics. While the observed upward shift in the age distribution of reported pertussis cases during the more recent major epidemics, e.g. [18] vs. [19], provides some indication to that effect, a better understanding of the role of different age groups, including older adolescents during pertussis outbreaks is needed.

Previously, we introduced a method for assessing the roles of different population groups during infectious disease outbreaks [20-22] and applied it to data from the 2012 pertussis epidemics in Minnesota and Wisconsin [16,23]. That inference method compares age groups in terms of their proportion among reported cases before vs. after the outbreak’s peak. Groups that play a more prominent role in perpetuating outbreaks due to either increased contact rates, or increased susceptibility to infection, or both, are overrepresented among cases of infection occurring during the ascent of the outbreak. Such groups experience a disproportionate depletion of the pool of susceptible individuals during the outbreak’s early stages and represent a relatively smaller proportion of all cases of infection in the population, as well as of reported cases during the outbreak’s later stages. Importantly, this comparison of the relative roles of different age groups does not depend on the differences in case reporting rates (proportion of cases of infection that are reported) in different age groups, as long as age-group-specific case reporting rates don’t change during the course of the outbreak [20]; potential changes in case-detection rates during the course of an epidemic are likely to bias the relative risk estimates towards the null (see Discussion). When applied to data from pertussis epidemics, that inference method suggested the prominent role of adolescents aged 11-14y during the 2012 pertussis outbreaks in Minnesota and Wisconsin [16,23].

In this paper, we apply the methodology in [20-22] to assess the relative roles of different age groups during the 2010 and 2014 pertussis outbreaks in California. Quantification of the relative role for an age group according to the methodology in [20-22] is related to the impact of vaccination of an individual in that age group at the start of an epidemic on reducing the epidemic’s initial growth rate/reproductive number (Supporting Information in each of [16,20,22]). Additionally, we examine the differences in the role of different age groups during the 2014 vs. the 2010 epidemic. This comparison is partly motivated by the possible rise in the importance of older adolescents during the more recent pertussis outbreaks, and the potential need for a booster dose, that is effective against infection and transmission, for older adolescents [6,10,15,16] with the aim of mitigating future pertussis epidemics in the community.

## Methods

### Data

We considered pertussis case reporting data between 2010-2015 collected by the California Department of Public Health. The analyses were restricted to the time period covering the duration of each outbreak. For the 2010-2011 epidemic, the analyses were restricted to the time period between week 1, 2010 and week 12, 2011. For the 2014 epidemic, the analyses were restricted to the period between weeks 1, 2014 and week 52, 2014. Data used in this study could be obtained from Dr. Kathleen Harriman, California Department of Public Health. We’ve used de-identified data on pertussis cases reported to the California Department of Public Health, and no informed consent from those individuals was sought.

We considered six regions in California that comprise the following counties:

Region 1: San Diego, Imperial, San Bernardino, Riverside

Region 2: Los Angeles, Orange, Ventura

Region 3: Kern, Tulare, Fresno, Madera, Merced, Mariposa, Tuolumne, Stanislaus, San Joaquin, Amador, Calaveras

Region 4: San Benito, Santa Clara, Santa Cruz, San Mateo, Alameda, Contra Costa, San Francisco, Marin, Solano, Napa, Sonoma

Region 5: Yolo, Sacramento, El Dorado, Yuba, Placer, Sutter, Butte, Nevada, Colusa, Glenn, Tehama, Shasta

Region 6: Santa Barbara, San Luis Obispo, Monterey

We note that the outbreak in each region may comprise multiple local outbreaks with peaks potentially occurring at different times than the regional peak. To mitigate the potential effect of this phenomenon on our inference method, the statistical analysis only included those regions where the incidence curves of reported pertussis cases had pronounced major peaks (Figure 1 and 2). The starting and ending weeks for each region was determined by the ascent and descent of the epidemic wave in that region. Section S1 of the Supporting Information gives further details on the selected regions, the starting and ending weeks for the major waves of the 2010-2011 and the 2014 pertussis epidemics in those regions, and the number of reported pertussis cases during those epidemic waves in the selected regions.

**Figure 1:**
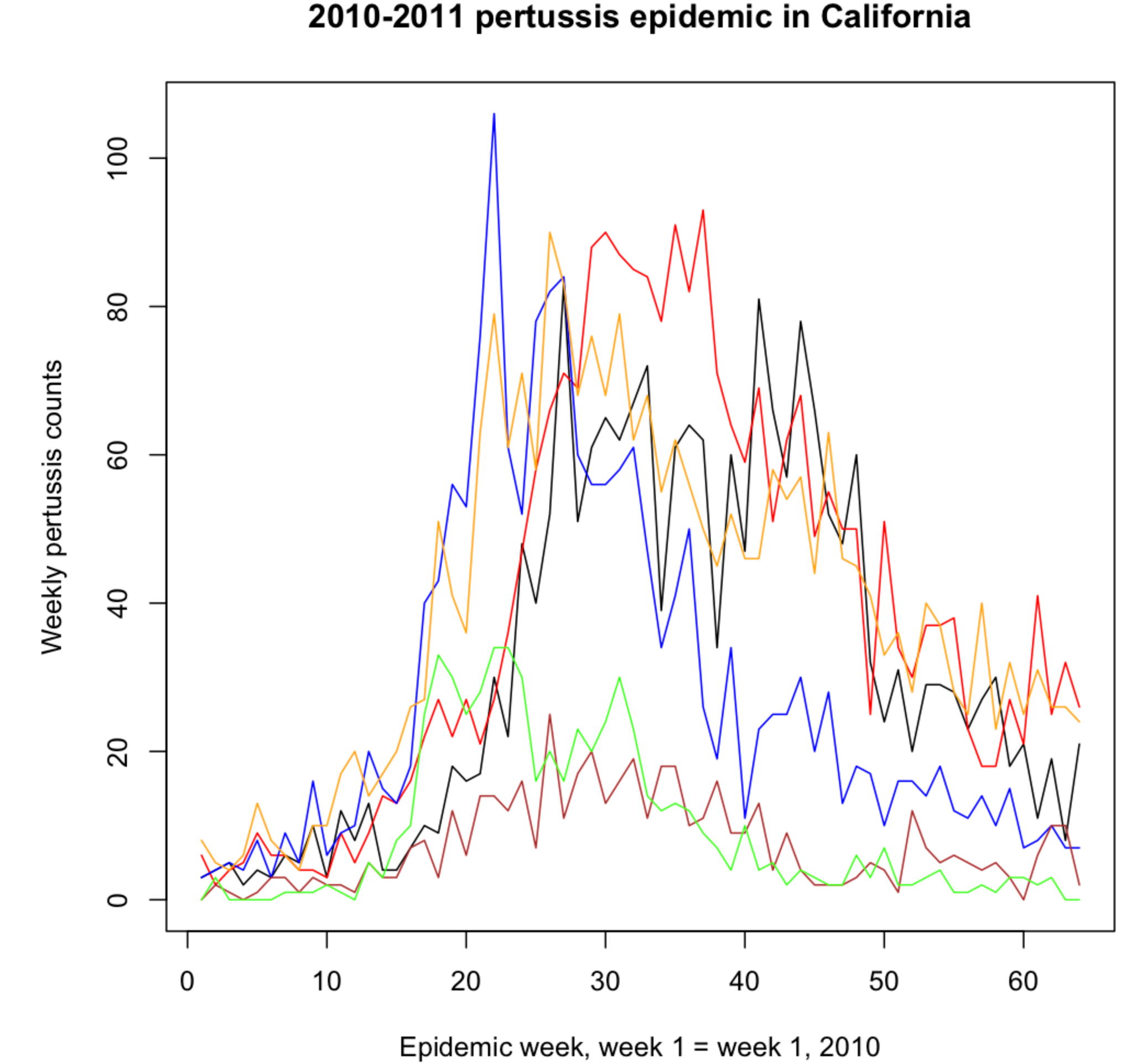
Weekly pertussis incidence (cumulative counts for all ages) for the reported pertussis cases in different California regions during the 2010-2011 epidemic

**Figure 2:**
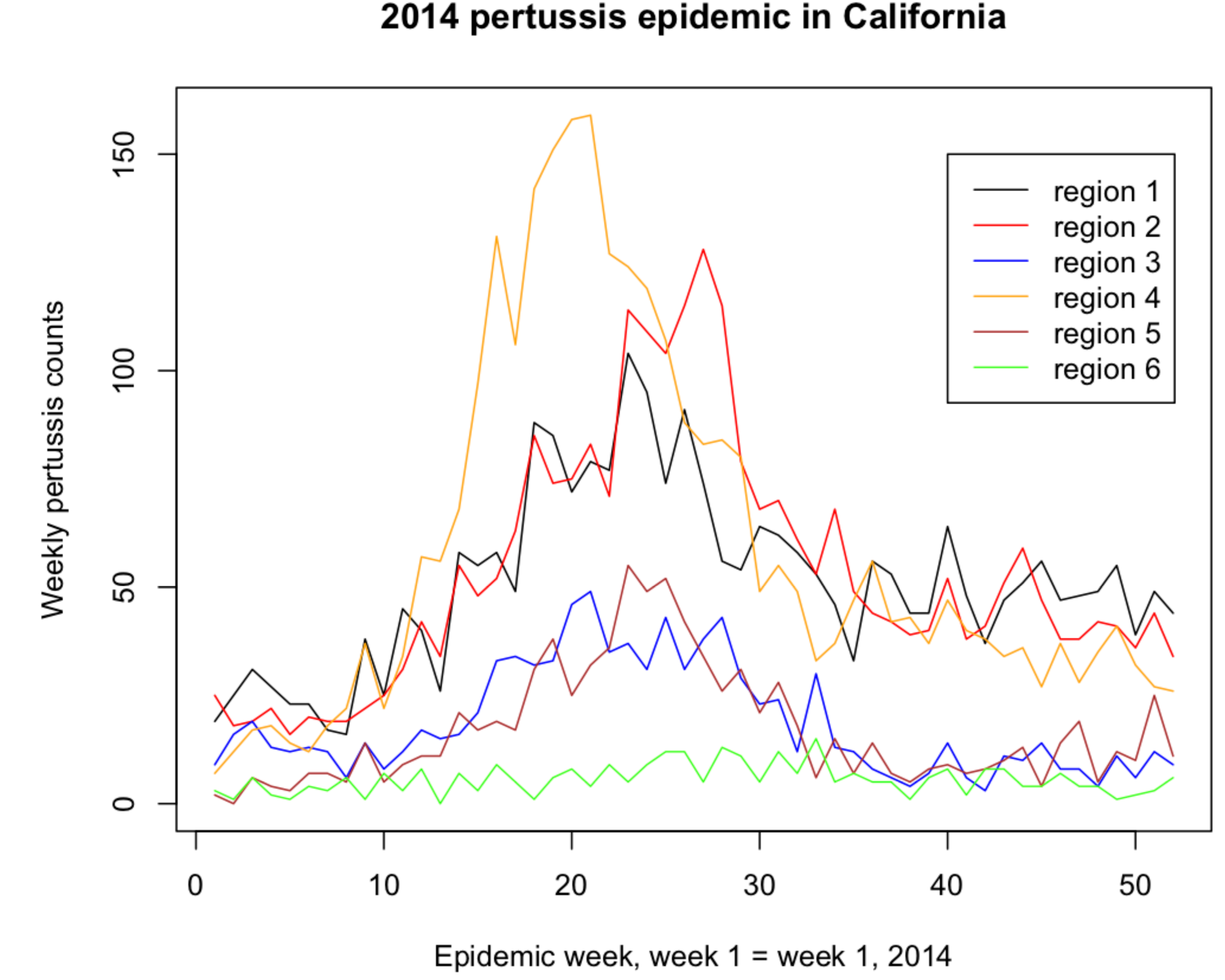
Weekly pertussis incidence (cumulative counts for all ages) for the reported pertussis cases in different California regions during the 2014 epidemic

### Statistical analysis

We categorized cases into ten age groups (at onset of illness), in years: (<1, 1-2, 3-4, 5-6, 7-8, 9-10, 11-13, 13-15, 14-15, 16-19, 20+). We used the region-specific outbreak peak times (namely weeks when the overall incidence of detected cases in a given region is highest) to determine whether reported cases occurred before or after the peak. The region-specific peak week *t* for reported cases (in all age groups) may not correspond to the peak week for the true incidence of pertussis infection in the community in that region because only a fraction of cases of pertussis infection are diagnosed and reported to the California Department of Public Health. To diminish the possibility of misclassification of cases as those occurring before or after the epidemic peak, we defined the regional before-the-peak period to be the period up to week *t*-*2* (inclusive), and the after-the-peak period to be period starting week *t* + *2*. Cases occurring during weeks *t* - 1 through *t* + 1 were excluded.

For the joint analysis for the included regions during each epidemic, for each age group *g*, cases occurring before the outbreak peak in each region were combined, with their total number denoted by *B*(*g*), and the same applies to cases occurring after the peak, with their number denoted by *A*(*g*). The estimated relative risk for each age group *g* is the ratio of the proportions of cases in the group *g* among all reported pertussis cases in the population before the peak and after the peak as in eq. 1 (here *h* in the sum runs over all age groups):

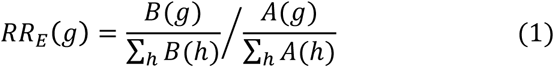

The observed numbers of reported cases *B*(*g*) and *A*(*g*) in the age group *g* before and after the peak are binomially distributed. Moreover, we assume that the numbers of reported cases are sufficiently high so that the logarithm In (*RR*(*g*)) of the relative risk in the age group *g* is approximately normally distributed ([24]). Under this approximation, the 95% confidence interval for *RR*(*g*) is *exp*(In(*RR*_*E*_(*g*))±1.96.*SE*), where In (*RR*_*E*_(*g*)) is estimated via eq. 1, and the standard error is

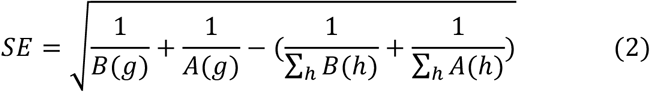

Estimation of the relative risks can also be performed in a Bayesian framework under the assumption that the proportion of cases of pertussis infection in each age group reported to the California Department of Public Health is small, which was found to be the case in other settings [25]. Details are given in section S3 of the Supporting Information.

## Results

Figures 1 and 2 plot the weekly numbers of reported pertussis cases in the six California regions described in the Methods for the 2010-2011 and the 2014 epidemics. We note that the regional incidence curves for the 2014 epidemic had more pronounced peaks compared to the regional incidence curves for the 2010-2011 epidemic. Figure 3 plots the incidence rates (per 100,000) for reported pertussis cased for the 2010-2011 and the 2014 epidemics in California for the age groups used in our analysis (Methods). Figure 3 suggests an upward shift in the age distribution for reported pertussis cases for the 2014 epidemic compared to the 2010-2011 epidemic.

**Figure 3:**
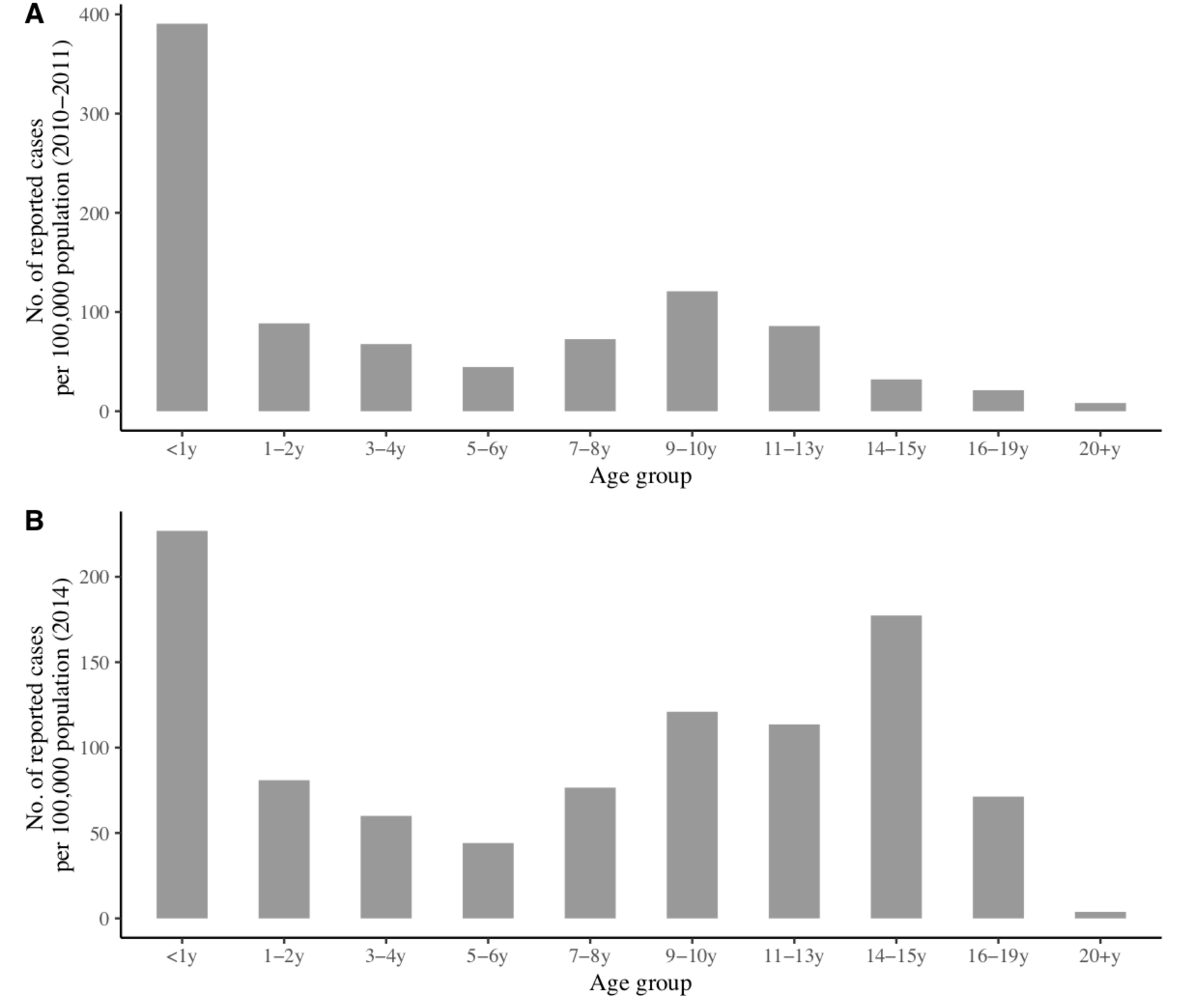
Age-specific pertussis incidence rates (per 100,000) for reported pertussis cases during the (A) 2010-2011 epidemic (week 1, 2010 through week 12, 2011), and (B) 2014 epidemic (weeks 1-52, 2014).

Table 1 shows the estimates of the relative risk (RR, eqns. 1 and 2) in the different age groups considered in our analyses for the 2010-2011 and the 2014 pertussis epidemics in California. For the 2010-11 epidemic, delineation of the groups with higher RR estimates showed modest evidence of a leading role for adolescents in transmission. The leading RR estimates were 1.26 (1.08,1.46) (adolescents aged 11-13y); 1.19 (1.01,1.4) (children aged 9-10y); 1.17 (0.86,1.59) (adolescents aged 14-15y); 1.12 (0.86,1.46) (aged adolescents 16-19y); and 1.1 (0.89,1.36) (children aged 7-8y). For the 2014 epidemic, a leading role for adolescents was clearer. The highest RR estimate belonged to adolescents aged 14-15y (RR=1.83, 95%CI(1.61,2.07)), followed by adolescents aged 16-19y (RR=1.41 (1.24,1.61)) and 11-13y (RR=1.26 (1.12,1.41)), with lower RR estimates in other age groups.

**Table 1:** RR estimates for different age groups during the 2010-11 and the 2014 pertussis epidemics in California (eq. 1)

| Age group | 2010-11 epidemic | 2014 epidemic |
| --- | --- | --- |
| <1y | 1.02 (0.9,1.14) | 0.73 (0.63,0.85) |
| 1-2y | 0.68 (0.56,0.83) | 0.54 (0.45,0.65) |
| 3-4y | 0.64 (0.51,0.82) | 0.5 (0.4,0.62) |
| 5-6y | 0.99 (0.76,1.31) | 0.46 (0.35,0.59) |
| 7-8y | 1.1 (0.89,1.36) | 0.88 (0.74,1.05) |
| 9-10y | 1.19 (1.01,1.4) | 0.98 (0.85,1.13) |
| 11-13y | **1.26 (1.08,1.46)** | 1.26 (1.12,1.41) |
| 14-15y | 1.17 (0.86,1.59) | **1.83 (1.61,2.07)** |
| 16-19y | 1.12 (0.86,1.46) | 1.41 (1.24,1.61) |
| 20+y | 0.95 (0.85,1.05) | 0.79 (0.67,0.92) |

Pairwise comparison of the role of individuals in different pairs of age groups during each epidemic is presented in section S2 of the Supporting Information. Additionally, estimates of the RR statistic in a Bayesian framework are given in section S3 of the Supporting Information, with the results being very similar to the results in Table 2.

## Discussion

A good deal of uncertainty exists about the roles of individuals in different age groups in propagating the recent pertussis outbreaks. Moreover, the observed upward shift in the age distribution of reported pertussis cases for the more recent outbreaks (e.g. [18] vs [19]) suggests the temporal changes in the role of different age groups during pertussis epidemics. In this paper, we use the previously developed methodology based on the relative risk (RR) statistic [20-22] to examine the roles of different age groups during the 2010 and the 2014 pertussis outbreaks in California, and compare those roles for the two epidemics. Our results suggest the prominence of adolescents aged 14-15 year during the 2014 pertussis epidemic, followed by adolescents aged 16-19 years and 11-13 years. We also note that the highest rates of detected cases during the 2014 epidemic in California outside the infant age group belonged to adolescents aged 14-15 years (Figure 3). For the 2010 pertussis epidemic in California, there was no strong differentiation in the role of different age groups based on the RR statistic, with the leading estimates of the RR statistic belonging to adolescents aged 11-13 years, followed closely by children aged 9-10 years, adolescents aged 14-15 years, 16-19 years, and children aged 7-8 years. Our earlier findings for the 2012 pertussis epidemics in Minnesota and Wisconsin [16,23] were consistent with the leading roles of adolescents aged 11-14 years during those outbreaks. The combination of the findings in this paper and the findings in [16,23] suggests that as time progressed, the prominence of older adolescents during pertussis outbreaks increased. This conclusion may be partly explained by the fact that adolescents of increasingly older age are covered entirely by the acellular pertussis vaccination series as the time from the introduction of acellular pertussis vaccines in different places grows, while the receipt of acellular vaccines alone is less protective against pertussis compared to receipt of some whole cell pertussis vaccination [4,5]. Correspondingly, older adolescents are expected to play the leading role during major future pertussis epidemics as well.

The findings about the prominence of older adolescents during the more recent pertussis outbreaks lead to questions about the protective effect of pertussis vaccination on individuals in those age groups, as well as the impact of vaccination on mitigating the spread of pertussis epidemics in the whole community. Under the current pertussis vaccination schedule [14], Tdap vaccine is usually administered around the age of 11 years, and its effectiveness against reportable pertussis disease wanes with time since vaccine administration [8,10,26]. Moreover, effectiveness of pertussis vaccines against infection and transmission to others may be lower compared to effectiveness against reportable pertussis disease [15]. While evidence about the herd immunity effects of DTaP is documented in the literature [17], the extent of the herd immunity effects of Tdap is less certain [16]. Further work is needed to better understand the effectiveness of acellular pertussis vaccines against infection and transmission to others. Such work should inform the potential impact and cost-effectiveness of modifications to the current pertussis vaccine schedule, such as vaccination of older adolescents, possibly with more efficacious vaccines against pertussis transmission than Tdap.

Our paper has some limitations. The relation between the RR statistic and the role played by individuals in a given age group during the outbreak is not entirely clear. The role of individuals in different age groups can be compared by comparing the effect of the distribution of a fixed quantity of a highly efficacious pertussis vaccine to members of one age group at a time at the beginning of an epidemic on the growth rate/reproductive number of the outbreak in the whole community. Our earlier work (see Supporting Information for each of the following papers [16,20,22]) had attempted to address this issue through simulations of transmission dynamics, finding a positive association between the RR statistic for a group and the per capita impact of vaccination in this group on the epidemic’s initial growth rate/reproductive number. However, this conclusion required certain assumptions the distribution of susceptibility to infection in different age groups ([22]). Additionally, the relative grading of different age groups according to the RR statistic does not depend on the differences in case reporting rates in different age groups, as long as case reporting rates don’t change during the course of the outbreak.

However, such changes are possible as awareness about the epidemic and the prominence of certain age groups may result in changes in testing and reporting practices during the course of the epidemic. For example, during the 2014 pertussis epidemic in California, the highest rates of detected cases outside the infant group were in adolescents aged 14-15y (Figure 3). If there was increase in awareness about pertussis incidence in those adolescents during the epidemic, case-detection rates in that age group are expected to rise as the epidemic progressed, and there would be more detected cases in adolescents aged 14-15y after the peak, biasing the RR estimate in that age group downward, while the RR estimate for adolescents aged 14-15y was higher than in other age groups (Table 1). More generally, changes in case detection rates due to awareness about the unusually high burden of pertussis disease in certain age groups are expected to bias the RR estimates towards the null, as suggested by the 2011 pertussis epidemic in England [27]. This novelty/change in awareness factor may be more true for the 2010 epidemic compared to the 2014 epidemic, where we were able to detect a strong signal despite potential biases towards the null.

In summary, our results suggest the shift in the role of different age groups during the 2014 pertussis epidemic in California compared to the 2010 epidemic, including the prominence of adolescents aged 14-15y during the 2014 epidemic. Additionally, older adolescents played a more prominent role during the 2014 pertussis epidemic in California compared to the 2012 pertussis outbreaks in Minnesota and Wisconsin [16,23]. Those findings are in agreement with the notion that as the time from the introduction of acellular pertussis vaccines in different places grows, older adolescents will be covered entirely by the acellular pertussis vaccination series, which is less protective against pertussis compared to receipt of some whole cell pertussis vaccination, particularly the priming dose [4,5]. Moreover, under the current pertussis vaccination schedule, the Tdap vaccine is usually administered around the age of 11 years, and its effectiveness wanes with time [8,10,26]. All of this suggests that older adolescents are expected to play a leading role in major future pertussis outbreaks as well. Pertussis activity in the US has decreased in the last few years, presumably at least partly due to the immunity imparted during high levels of *B. pertussis* circulation between 2010-2014 [1]. Pertussis activity is expected to increase with time as the population immunity wanes, and there is a need to examine various questions related to pertussis vaccination policies to better manage future outbreaks. One of those questions is the potential impact of vaccinating older adolescents with the aim of mitigating the spread of pertussis in the whole community, including the disease burden in infants. Such vaccination strategies require pertussis vaccines with high effectiveness in preventing pertussis infection and transmission to others, rather than just pertussis disease [15]. There is uncertainty about the effectiveness of Tdap in preventing pertussis infection and transmission to others (rather than effectiveness against detectable symptomatic pertussis disease in vaccine receipients), as well as the temporal waning of such effectiveness. Further work is needed in this direction, including the study of the potential effect of the deployment of more efficacious vaccines against pertussis infection in adolescents as well as the impact of vaccination of older adolescents on the spread of pertussis in the whole community, including disease rates in infants.

## Acknowledgement

We thank Sarah New, Kathleen Winter, and Kathleen Harriman of the California Department of Public Health, Immunization Branch for providing the data used in the analyses, and for helpful suggestions on the manuscript.

## Supporting Information for On the role of different age groups during pertussis epidemics in California, 2014 and 2010

### Section S1

Selection of regions and starting and ending weeks for the 2010 and 2014 epidemics

Table S1 shows the starting and ending weeks for the major waves of the 2010 and the 2014 pertussis epidemics in 6 different regions in California. Each region comprises a number of counties whose names are listed in Table S1. The selection of counties that constitute different regions was largely guided by the list of economic regions of California[1], with some exceptions. In particular, we’ve grouped the San Diego, Imperial, San Bernardino, and Riverside counties into one region, excluded several counties east of the Sierra Nevada and in the northern part of the state, and included the Mariposa, Tuolumne, Calaveras, and Amador counties into region 3. The starting and ending weeks were selected by inspection of Figures 1 and 2 in the main text to be the weeks when the ascent of the first major wave of the epidemic began, and the descent of that wave ended, correspondingly. Regions that did not have a pronounced epidemic peak (as suggested by visual inspection) were not included in the relative risk (RR) analysis as such regions may comprise further sub-regions with asynchronous epidemics, leading to a misclassification of a large number of cases as being before or after the peak of the epidemic.

**Table S1:** Starting and ending weeks for the major waves of the 2010 and the 2014 pertussis epidemics (calendar weeks for the corresponding year) in different regions in California. “X” indicates that the region wasn’t selected for the relative risk analysis.

| Region | 2010 |  | 2014 |  |
| --- | --- | --- | --- | --- |
|  | Starting week | Ending week | Starting week | Ending week |
| 1. San Diego, Imperial, San Bernardino, Riverside | X | X | 8 | 35 |
| 2. Los Angeles, Orange, Ventura | 10 | 49 | 5 | 38 |
| 3. Kern, Tulare, Fresno, Madera, Merced, Mariposa, Tuolumne, Stanislaus, San Joaquin, Amador, Calaveras | 6 | 40 | 10 | 38 |
| 4. San Benito, Santa Clara, Santa Cruz, San Mateo, Alameda, Contra Costa, San Francisco, Marin, Solano, Napa, Sonoma | 8 | 38 | 6 | 33 |
| 5. Yolo, Sacramento, El Dorado, Yuba, Placer, Sutter, Butte, Nevada, Colusa, Glenn, Tehama, Shasta | X | X | 5 | 33 |
| 6. Santa Barbara, San Luis Obispo, Monterey | 3 | 39 | X | X |

The 2010 epidemic had 5,422 reported cases in the selected regions between the starting and ending weeks of the epidemic in those regions; the 2014 epidemic had 7,440 reported cases in the selected regions between the starting and ending weeks of the epidemic in those regions.

### Section S2

Odds ratios for different pairs of age groups

The relative risk (RR) statistic (eq. 1 in the main text) allows for the simultaneous comparison of all age groups in terms of the relative depletion of susceptible individuals in those age groups before the epidemic peak. For a pairwise comparison of different age groups *ag*_1_, *ag*_*2*_, we consider reported pertussis cases that were either in *ag*_1,_or *ag*_*2*_, and evaluate the odds ratio *OR*(*ag*_1_, *ag*_*2*_) for being in *ag*_1_ vs *ag*_2_ for cases before vs after the epidemic peak. The estimate *OR*(*ag*_1_, *ag*_*2*_)>1means that the proportion of cases in *ag*_1_ among all cases in the two age groups had decreased after the peak, suggesting a higher depletion of susceptible individuals in *ag*_1_ compared to *ag*_2_ before the epidemic peak; the estimate *OR*(*ag*_1_, *ag*_*2*_)< 1 suggests a higher depletion of susceptible individuals in *ag*_2_ compared to *ag*_1_ by the time of the epidemic peak. We compute the odds ratio *OR*(*ag*_1_, *ag*_2_) using a logistic regression model, adjusting for whether the case was Hispanic, and whether the case was white. We note that the unadjusted odds ratio for a pair of age groups *ag*_1_,*ag*_2_ is simply the ratio of the relative risks *RR* (*ag*_1_) and *RR* (*ag*_2_) in those age groups[2]:

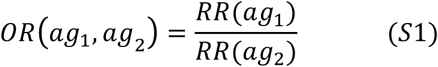

Here *RR*(*ag*_1_) and *RR*(*ag*_2_) are given by eq. 1 in the main text, and eq. S1 can be established simply by plugging the expressions for *RR*(*ag*_1_) and *RR*(*ag*_2_) in eq. 1 in the main text into the ratio 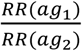.

Tables S2 and S3 present the estimates of the odds ratios *OR*(*ag*_1_, *ag*_2_) for different pairs of age groups during the 2010 and the 2014 epidemics in California.

Table S2 suggests a lower depletion of susceptible individuals in children aged 2-4y compared to all other age groups during the 2010 epidemic, and a higher depletion of susceptible individuals in children aged 9-13y compared to adults aged over 20y during the 2010 epidemic.

Table S3 suggests, among other things, a greater depletion of susceptible individuals in adolescents aged 14-15y compared to adolescents aged 11-13y and 16-19y during the 2014 epidemic; a higher depletion of susceptible individuals in adolescents aged 11-19y compared to other age groups during the 2014 epidemic; a higher depletion of susceptible individuals in children aged 7-10y compared to children aged 2-6y during the 2014 epidemic; and a higher depletion of susceptible individuals in children aged 9-10y compared to infants aged <1y and adults aged over 20y during the 2014 epidemic.

**Table S2:**
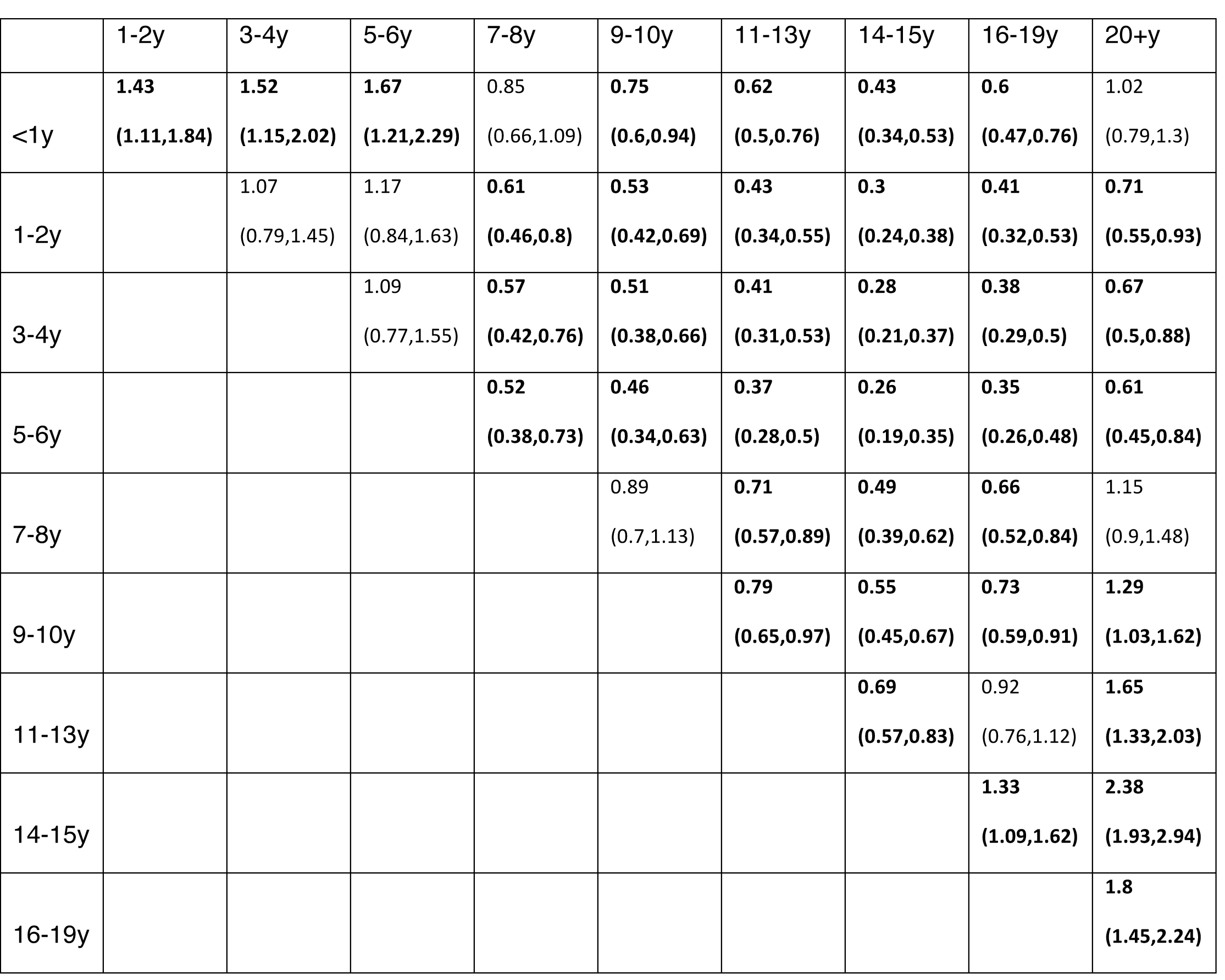
Odds ratios for being before vs after the peak of the 2010 pertussis epidemic in California for reported pertussis cases in members of one age group vs another for different pairs of age groups. Significant estimates are marked in bold.

**Table S3:**
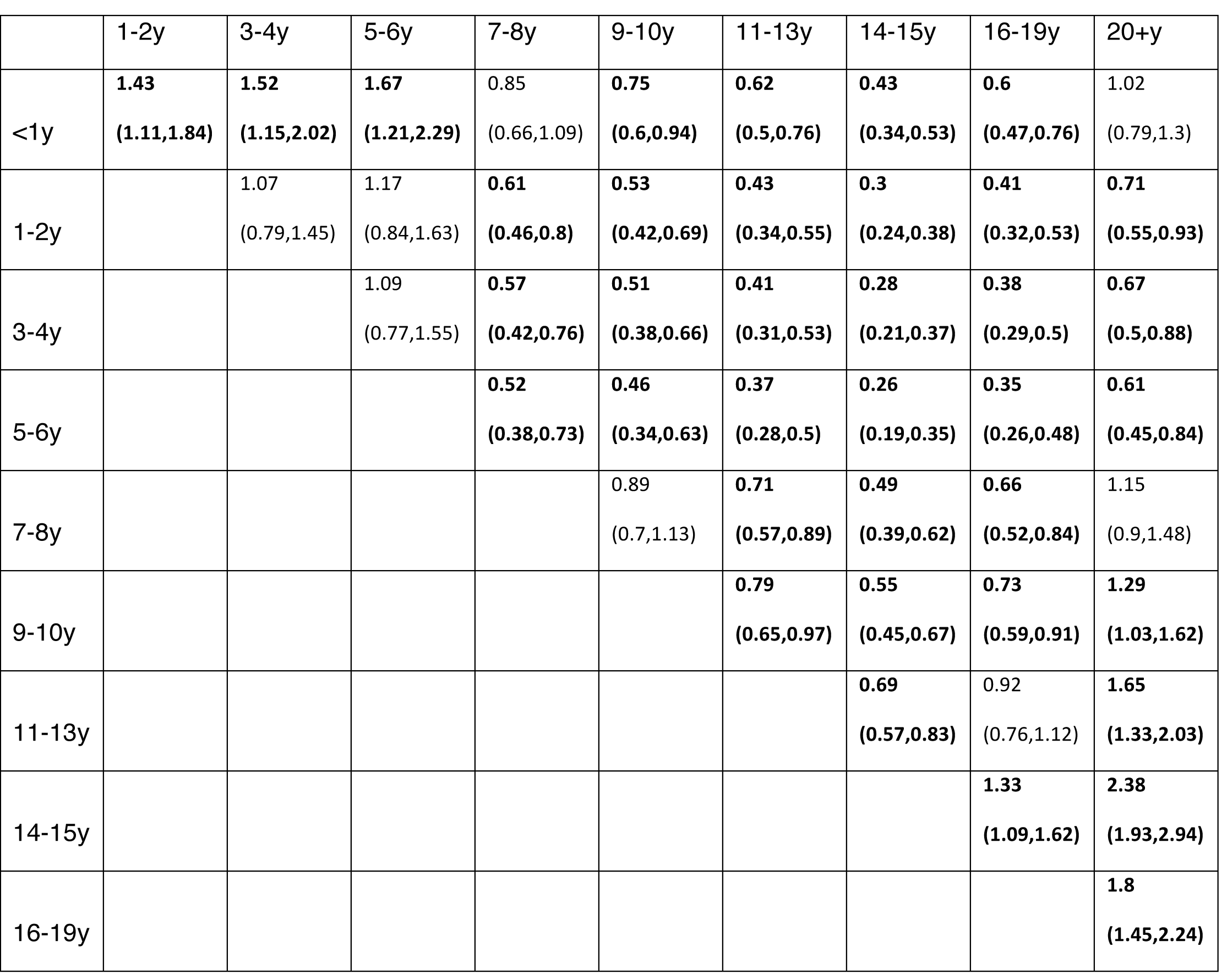
Odds ratios for being before vs after the peak of the 2014 pertussis epidemic in California for reported pertussis cases in members of one age group vs another for different pairs of age groups. Significant estimates are marked in bold.

### Section S3

RR estimation in a Bayesian framework

Derivation of the confidence bounds for the RR estimates using eq. 2 in the main text relies on a normal approximation that is valid for large sample sizes. Here, we derive estimates/credible intervals for the RR statistic in different age groups in a Bayesian framework.

For each epidemic and each age group *g*, cases occurring before the outbreak peak in each region included in the analysis were combined, with their total number denoted by *B*(*g*), and the same applies to cases occurring after the peak, with their number denoted by *A*(*g*). The expression for the relative risk for the age group *g* is the ratio of the proportions of cases in the group among all reported *g* cases in the population before the peak and after the peak as in eq. 1 in the main text (here *h* in the sum runs over all age groups).

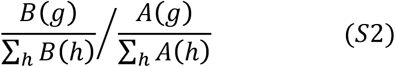

The estimates and confidence bounds for relative risks in each group can be obtained in a Bayesian framework based on the observations {*B*(*h*), *A*(*h*)} following the methodology in[2]. Briefly, if the number of cases of pertussis infection in each age group is large and case-reporting rates (proportion of cases of pertussis infection in a given age group that are reported to the California Department of Public Health) are low[3], the observed numbers of reported cases in each age group before and after the peak are Poisson distributed (with unknown true Poisson parameters). Posterior samples for the Poisson parameters (with a flat prior) corresponding to the observed counts {*B*(*h*), *A*(*h*)} are generated; for each i=1,..,100000, the corresponding parameters {*B*^*i*^(*h*), *A* ^*i*^(*h*)} are plugged into equation (S2) to generate an estimate *RR*^*i*^*(g)* for the relative risk in the age group *g*. The mean and the credible interval for the sample *(RR*^*i*^*(g)*) (i=1,..,100000) are then extracted. Table S4 exhibits the RR estimates in different age groups for the 2010 and the 2014 pertussis epidemics in California. Those estimates are very similar to the estimates in Table 1 in the main text.

**Table S4:** RR estimates for different age groups during the 2010 and the 2014 pertussis epidemics in California in a Bayesian framework

| Age group | 2010-11 epidemic | 2014 epidemic |
| --- | --- | --- |
| <1y | 1.02 (0.9,1.14) | 0.74 (0.63,0.85) |
| 1-2y | 0.69 (0.56,0.83) | 0.55 (0.45,0.65) |
| 3-4y | 0.65 (0.51,0.81) | 0.5 (0.4,0.62) |
| 5-6y | 1 (0.76,1.3) | 0.46 (0.35,0.59) |
| 7-8y | 1.11 (0.89,1.36) | 0.88 (0.74,1.05) |
| 9-10y | 1.19 (1.01,1.4) | 0.99 (0.85,1.13) |
| 11-13y | **1.26 (1.08,1.46)** | 1.26 (1.12,1.41) |
| 14-15y | 1.18 (0.86,1.58) | **1.83 (1.61,2.07)** |
| 16-19y | 1.13 (0.86,1.46) | 1.41 (1.24,1.61) |
| 20+y | 0.95 (0.85,1.05) | 0.79 (0.67,0.92) |

